# Chromoanagenesis in the *asy1* meiotic mutant of *Arabidopsis*

**DOI:** 10.1101/2022.04.27.489737

**Authors:** Weier Guo, Luca Comai, Isabelle M. Henry

## Abstract

Chromoanagenesis is a catastrophic event that involves localized chromosomal shattering and reorganization. In this study, we report a case of chromoanagenesis resulting from defective meiosis in the MEIOTIC ASYNAPTIC MUTANT 1 (*asy1*) background in *Arabidopsis thaliana*. We provide a detailed characterization of the genomic structure of this individual with a severely shattered segment of chromosome 1. We reveal more than 300 candidate novel DNA junctions in the affected region, confirming that *asy1*-related defective meiosis is a potential trigger for chromoanagenesis. This is the first example of chromoanagenesis associated with female meiosis and indicates the potential for genome evolution during oogenesis.

## Background

Complex chromosomal rearrangements (CCRs) refer to genomic structure variation that involve at least three double strand DNA breaks among two or more chromosomes [1]. These changes can cause the truncation, relocation, or copy number variation of multiple genes or gene regulatory elements, which can subsequently lead to dramatic phenotypic changes [2]. Chromoanagenesis, caused by a single catastrophic genome restructuring event, and diagnosed by the presence of tens to hundreds of copy number variations (CNVs) on a single chromosome, has been identified in many systems in the last decade [3–10]. It can be associated with multiple types of human cancer [9, 11], or with transgenic modifications used for plant genetic engineering [7, 10, 12, 13]. The origin, mechanism and potential effect of chromoanagenesis are just starting to be deciphered.

MEIOTIC ASYNAPTIC MUTANT 1 (ASY1), the arabidopsis homolog of the yeast chromosome axis component HOP1, plays an important role in meiotic recombination by regulating crossover assurance and interference [14–16]. First observed in transgenic Arabidopsis mutants exhibiting reduced synapsis [14, 17], aneuploidy in the progeny of ASY1 mutants suggested that ASY1 mutation can also result in genome instability [18, 19].

Here, we report a case of chromoanagenesis resulting from defective meiosis in the *asy1* mutant background in *Arabidopsis thaliana*. Specifically, a homozygous *asy1* mutant was crossed as a female to a wild-type male, and aneuploids were observed in the progeny. Detailed characterization of the genome of one of these aneuploid individuals detected a severely shattered segment of chromosome 1, which was reminiscent of the consequences of chromoanagenesis. Our analyses identified more than 300 potential novel DNA junctions in this region, suggesting that defective *asy1* is a potential process triggering chromoanagenesis.

## Results and Discussion

A recent study [20] demonstrated that the genome of one offspring from a cross between a Col-0/Ler-1 hybrid *asy1* mutants (asy1^*Col-0*^ x asy1^*Ler-1*^, female) and a wild-type Col-0 (male), carries drastic genomic rearrangement. These rearrangements resemble the consequence of chromoanagenesis. Among the population of 176 individuals, one line exhibited multiple CNVs, on chromosome 1, all clustered within the first half of the chromosome (from 1 to 16.1Mb) (Fig.1A-C).

**Figure 1.**
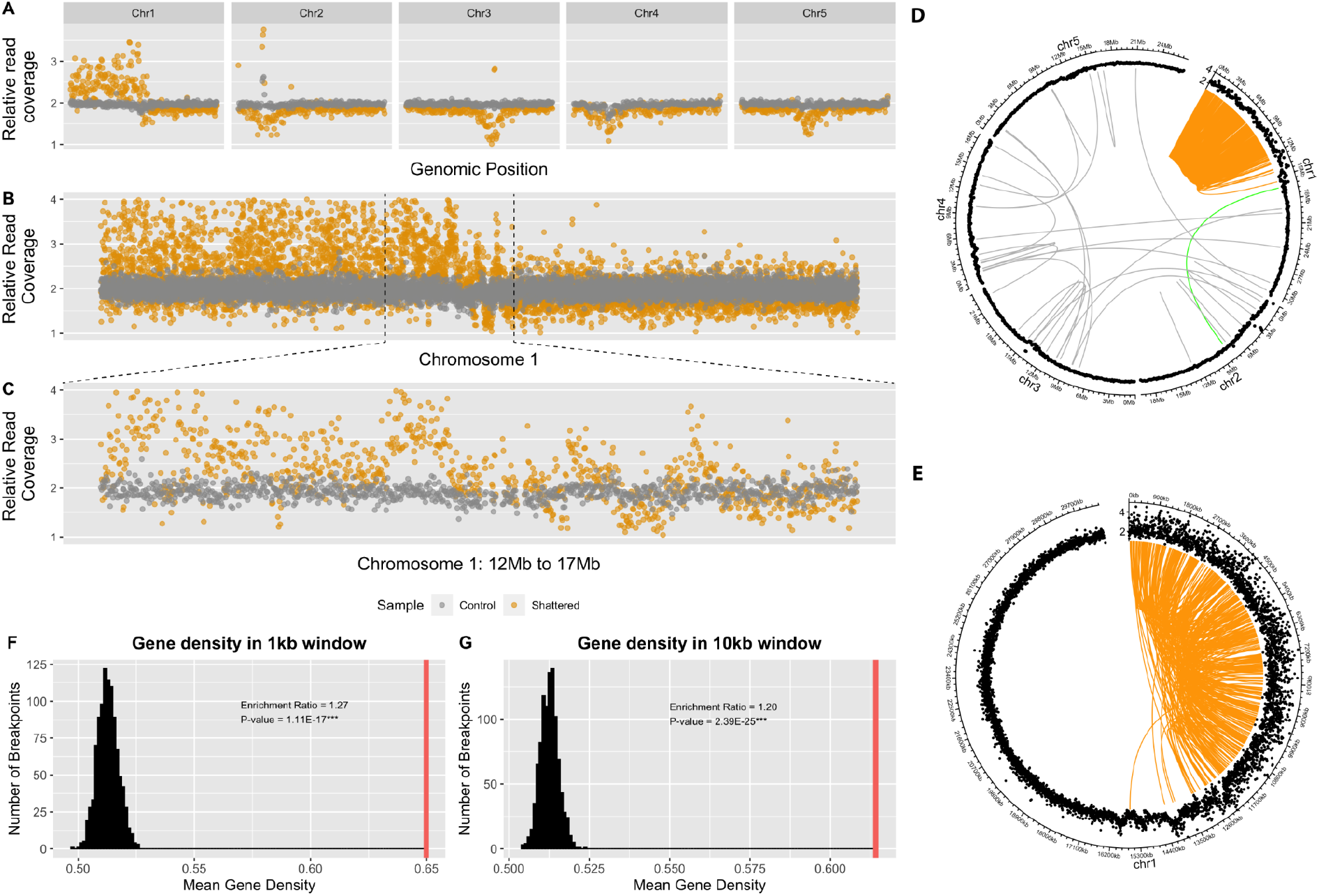
Characteristics of the genomic region with extreme dosage variations in the progeny of *asy1* mutant *A.thaliana*. (A-C) Extremely dense copy number variations on chromosome 1. Each dot represents the normalized read coverage in a bin set along the genome. (A) Relative read coverage across the whole genome (100kb bins). (B) Close-up of chromosome 1 (5kb bins). (C) Further close-up on the region of chromosome 1 that displays dense CNVs (5kb bins). (D,E) Candidate breakpoints were highly enriched over the regions of chromosome 1 exhibiting clustered CNVs. The distribution of candidate DNA junctions on all chromosomes of the arabidopsis genome (D) and just Chromosome 1 (E) are shown with circos plots. The outermost layer indicates chromosomes. The next layer indicates relative read coverage, with 100kb bins (D) and 5kb bins (E). The center arcs represent the locations of breakpoints pairs of the candidate DNA junctions identified. They are colored in orange if both breakpoints are located within the CNV cluster, in green if only one breakpoint falls within the CNV cluster, and in gray if both breakpoints fall elsewhere in the genome. The CNV cluster represents the first 16.1Mb of Chromosome 1. (F,G) Candidate novel breakpoints significantly occurred in gene rich regions. The black distribution represents the mean gene density of 1,000 pseudo breakpoints datasets (each dataset contains 1,000 pseudo breakpoints), and the red vertical line represents the mean gene density of observed candidate novel breakpoints. Enrichment ratios represent the comparison between the mean gene density of candidate novel breakpoints and the mean gene density of pseudo breakpoints. Both 1kb (F) and 10kb (G) windows show enrichment ratio > 1, suggesting the higher gene density around candidate novel breakpoints than the genome average (p-values both < 0.001).

**Figure 2.**
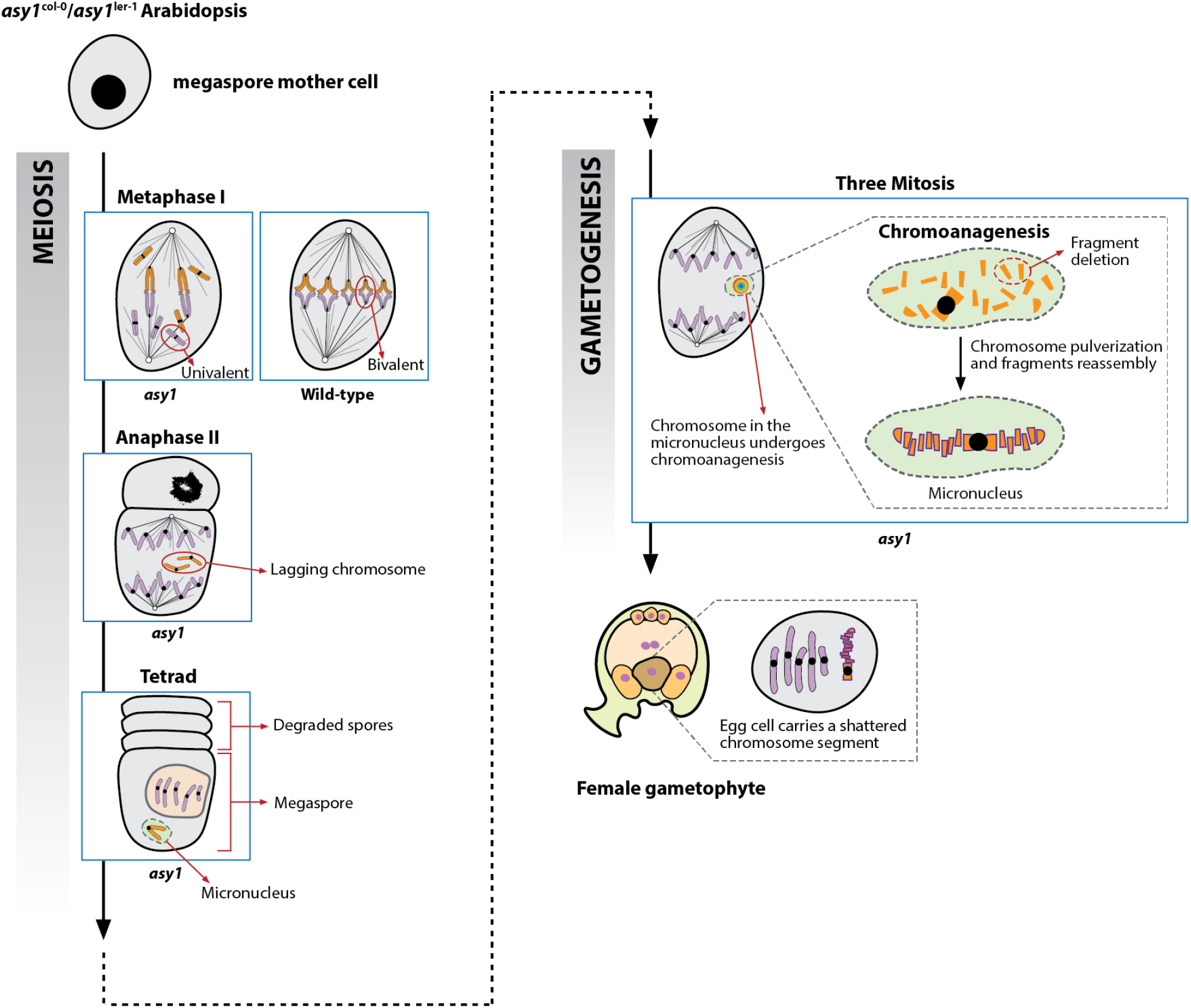
Proposed mechanism for chromoanagenesis in the *asy1* homozygous mutant. The megaspore mother cell from *asy1* homozygous mutant exhibits chromosome mis-segregation during female meiosis. Specifically, the *asy1* mutation results in the formation of univalents at metaphase I, which leads to unbalanced chromosome segregation. During meiosis II, the mis-segregated chromosome lags, and is incorporated into a micronucleus. In the following three mitosis during gametogenesis, the chromosome within micronucleus is unable to synchronize with the mitotic division of the main nucleus, and undergoes pulverization and restructuring, resulting in a chromosome with clustered structural variation. This shattered chromosome can be transmitted to the progeny if it is partitioned into the egg cell after micronucleus disassembly.

To confirm the occurrence of extreme chromosomal rearrangement, we searched for novel DNA junctions expected at the sites of chromosomal fragments reassembly. Specifically, we searched for Illumina sequencing reads that mapped to two distant genomic locations (>2,000bp), indicating that two regions expected to be distant from each other in the reference genome are next to each other in the rearranged chromosome. We also expect that these novel DNA junctions are unique to the genome of this particular individual and not present in its siblings. Based on these criteria, we identified more than 600 loci within this shattered region, representing over 300 candidate novel DNA junctions (Fig.1D, 1E). For about 91% (306 out of 336) of these candidate junctions, both breakpoints fall within the shattered region on chromosome 1 (Fig.1E). This frequency of one breakpoint every 25 kb on average is much higher than those observed following chromoanagenesis in other plant systems. Specifically, the frequency of breakpoints was 1 / 400 kb in chromoanagenetic individuals originated from haploid induction crosses in *A. thaliana* [7], and 1 breakpoint per 250 kb for the chromoanagenesis events observed in the progeny of gamma irradiated poplar pollen grains [10].

To characterize the properties of these novel DNA junctions, we investigated the DNA sequence context among candidate breakpoint loci. Two window sizes, 1kb and 10kb, were used to calculate the enrichment ratio of gene space around the candidate breakpoint loci. Statistical analysis suggested that breakpoint loci were significantly associated with gene-rich regions for both 1kb and 10kb window sizes (p-value < 0.001) (Fig.1F, 1G). These results are consistent with previously documented chromoanagenesis events in plants, which also exhibited higher than expected frequency of breakpoints occurrence in genic regions [7, 10]. In addition to gene density, we also characterized the potential enrichment of other genomic features, including chromatin states, transposable elements, or replication origins. The results suggest that breakpoint loci occurred more often in accessible chromatin regions, such as near transcription start sites, while they were significantly depleted in heterochromatin regions (Table 1).

**Table 1.** Enrichment ratio of novel breakpoints on other genomic features. ***: p-value < 0.001; **: p-value < 0.01; *: p-value < 0.0.5.

| Genomic features |  | 1kb window |  | Description |
| --- | --- | --- | --- | --- |
|  |  | Enrichment Ratio | P-value |  |
| Chromatin states | Chromatin state 1 | 1.38 | <0.001*** | TSS, promoter, 5' UTR |
|  | Chromatin state 2 | 1.21 | 0.06 | intergenic regions with proximal promoter elements |
|  | Chromatin state 3 | 1.5 | <0.001*** | transcription elongation signature |
|  | Chromatin state 4 | 0.86 | 0.05 | intergenic regions with distal promoter elements |
|  | Chromatin state 5 | 0.91 | 0.32 | Polycomb-regulated chromatin, intergenic region |
|  | Chromatin state 6 | 1.17 | 0.14 | gene bodies, intragenic state |
|  | Chromatin state 7 | 1.26 | 0.03* | intragenic region, 55.6% coding sequence, 34.3% introns |
|  | Chromatin state 8 | 0.58 | <0.001*** | AT-rich heterochromatin, high TE, low TE genes |
|  | Chromatin state 9 | 0.23 | <0.001*** | GC-rich heterochromatin, high TE and TE genes |
| DNase I hypersensitive sites |  | 1.28 | <0.001*** |  |
| Gene |  | 1.27 | <0.001*** |  |
| mRNA |  | 1.14 | <0.001*** |  |
| Replication origin |  | 1.4 | 0.03* |  |
| Transposable element |  | 0.44 | <0.001*** |  |

Notably, the vast majority of the previously identified chromoanagenesis events in animals [21–25], and all of the characterized events in plants [7, 10], have been associated with mitosis, usually during early embryo development [7, 26], or male gametogenesis [10]. Only a few studies have reported that chromoanagenesis can be correlated with meiotic divisions. Specifically, in human germ cells, chromoanagenesis has been demonstrated to occur during the meiotic divisions of spermatogenic cells and spermiogenesis [21, 27]. Extreme chromosomal rearrangements are also expected to occur following defects in female meiosis [28], but no case has been observed so far.

In this study, chromoanagenesis was detected in the offspring of an *asy1* homozygous mutant. Since ASY1 is involved in crossover assurance and interference, its deficiency resulted in altered recombination patterns and unbalanced chromosome segregation during meiosis [16]. Cytological evidence has shown the presence of unequal chromosome segregation during microsporogenesis in *asy1* mutants [17, 18, 29, 30]. Cytological analysis of female sporogenesis in these plants also documented abnormal chromosome pairing and uneven chromosome segregation [31, 32]. Moreover, haplotype analysis of the shattered chromosome 1 is consistent with meiosis I mis-segregation. These results suggest that unbalanced segregation of chromosomes occurs during megasporogenesis and may induce micronuclei formation. Micronuclei have been observed during male sporogenesis in *asy1* mutants with Col/Ler background [20]. It is thus possible that female sporogenesis also undergoes unbalanced chromosome segregation and subsequently produces micronuclei carrying a chromosome laggard. Fragmentation and reorganization of the chromosome entrapped within micronuclei subsequently creates the shattered chromosome. Together, our results provide the first example of chromoanagenesis triggered during female meiosis.

## Conclusions

We describe a case of chromoanagenesis that is remarkable by the high frequency of new DNA junctions produced, and because it results from asynapsis during female meiosis. The event demonstrates the potential for karyotypic innovation in connection to oogenesis.

## Material and Methods

DNA from the Arabidopsis line exhibiting multiple CNVs was prepared for deep sequencing as follows. The genomic DNA was extracted from the leaf tissue, and prepared for Illumina short-read sequencing as previously described [33]. Demultiplexing and quality filtering was performed using a custom Python script (https://comailab.org/data-and-method/barcoded-data-preparation-tools-documentation/). Reads were mapped to the TAIR10 reference genome using BWA [34]. The output files (.sam files) were used for the subsequent analyses. Two controls were generated by pooling the low-sequencing read data from multiple wild-type Arabidopsis lines generated from a similar cross (Ler-1/Col-0 x Col-0) (Supplementary File 1) to obtain two control files of similar coverage as the target sample.

Dosage variation along non-overlapping consecutive bins spanning the entire genome was documented as previously described [7, 33]. Bin coverage was normalized to the corresponding bin in a diploid control individual for normalization, by using a customized Python script (https://github.com/Comai-Lab/bin-by-sam). The expected relative read coverage of a diploid individual is expected to be close to 2, while values close to 1 and 3 represent deletion and duplication, respectively.

Candidate novel DNA junctions were identified as described previously [10]. Specifically, we searched for sequencing reads that span two genomic locations originally located at distant positions (>2000 bp apart, or on different chromosomes), and that appear uniquely in the target arabidopsis line but not either of the two control samples. A custom Python script (https://github.com/guoweier/Poplar_Chromoanagenesis) was used to identify the exact genomic locations of the two breakpoints for each novel junction. Potential false positives were discarded based on a coverage threshold calculated as previously described [10]. To analyze the genomic features surrounding the candidate breakpoints, the frequency of gene space surrounding them was compared to the frequency of pseudo breakpoints randomly selected along the Arabidopsis genome. The annotation files of various genomic features of *Arabidopsis thaliana* (TAIR10) was acquired from the GitHub repository (https://github.com/KorfLab/FRAG_project) associated with the breakpoint analysis previously performed on aneuploid Arabidopsis [7]. Statistical analysis was performed as previously described [10].

## Data availability

The sequences reported in this paper have been deposited in the National Center for Biotechnology Information BioProject database (BioProject ID: PRJNA723952).

## Authors’ contributions

WG analyzed and interpreted the data, and drafted the manuscript. LC and IMH conceptualized the project, and reviewed and edited the manuscript. IMH managed the project. All authors read and approved the final manuscript.

## Acknowledgments

We thank Dr Arp Schnittger for providing DNA from the chromothriptic individual. We thank the UC Davis DNA Technologies core for sequencing assistance.

## Supplementary Files

Supplementary File 1. List of the wild-type Arabidopsis lines used for generating two controls.

## References

1. Pellestor F, Anahory T, Lefort G, Puechberty J, Liehr T, Hédon B, Sarda P (2011) Complex chromosomal rearrangements: origin and meiotic behavior. Hum Reprod Update 17:476–494

2. Poot M, Haaf T (2015) Mechanisms of Origin, Phenotypic Effects and Diagnostic Implications of Complex Chromosome Rearrangements. Mol Syndromol 6:110–134

3. Stephens PJ, Greenman CD, Fu B, et al (2011) Massive genomic rearrangement acquired in a single catastrophic event during cancer development. Cell 144:27–40

4. Liu P, Erez A, Nagamani SCS, et al (2011) Chromosome catastrophes involve replication mechanisms generating complex genomic rearrangements. Cell 146:889–903

5. Baca SC, Prandi D, Lawrence MS, et al (2013) Punctuated evolution of prostate cancer genomes. Cell 153:666–677

6. Anand RP, Tsaponina O, Greenwell PW, Lee C-S, Du W, Petes TD, Haber JE (2014) Chromosome rearrangements via template switching between diverged repeated sequences. Genes Dev 28:2394–2406

7. Tan EH, Henry IM, Ravi M, Bradnam KR, Mandakova T, Marimuthu MPA, Korf I, Lysak MA, Comai L, Chan SWL (2015) Catastrophic chromosomal restructuring during genome elimination in plants. Elife 4:1–16

8. Blanc-Mathieu R, Krasovec M, Hebrard M, et al (2017) Population genomics of picophytoplankton unveils novel chromosome hypervariability. Sci Adv 3:e1700239

9. Cortés-Ciriano I, Lee JJK, Xi R, et al (2020) Comprehensive analysis of chromothripsis in 2,658 human cancers using whole-genome sequencing. Nat Genet 52:331–341

10. Guo W, Comai L, Henry IM (2021) Chromoanagenesis from radiation-induced genome damage in Populus. PLoS Genet 17:e1009735

11. Koltsova AS, Pendina AA, Efimova OA, Chiryaeva OG, Kuznetzova TV, Baranov VS (2019) On the complexity of mechanisms and consequences of chromothripsis: An update. Front Genet 10:393

12. Liu J, Nannas NJ, Fu F-F, Shi J, Aspinwall B, Parrott WA, Dawe RK (2019) Genome-Scale Sequence Disruption Following Biolistic Transformation in Rice and Maize. Plant Cell 31:368–383

13. Fossi M, Amundson K, Kuppu S, Britt A, Comai L (2019) Regeneration of Solanum tuberosum Plants from Protoplasts Induces Widespread Genome Instability. Plant Physiol 180:78–86

14. Caryl AP, Armstrong SJ, Jones GH, Franklin FC (2000) A homologue of the yeast HOP1 gene is inactivated in the Arabidopsis meiotic mutant asy1. Chromosoma 109:62–71

15. Sanchez-Moran E, Osman K, Higgins JD, Pradillo M, Cuñado N, Jones GH, Franklin FCH (2008) ASY1 coordinates early events in the plant meiotic recombination pathway. Cytogenet Genome Res 120:302–312

16. Lambing C, Kuo PC, Tock AJ, Topp SD, Henderson IR (2020) ASY1 acts as a dosage-dependent antagonist of telomere-led recombination and mediates crossover interference in Arabidopsis. Proc Natl Acad Sci U S A 117:13647–13658

17. Ross KJ, Fransz P, Armstrong SJ, Vizir I, Mulligan B, Franklin FC, Jones GH (1997) Cytological characterization of four meiotic mutants of Arabidopsis isolated from T-DNA-transformed lines. Chromosome Res 5:551–559

18. Wei F, Zhang G-S (2010) Meiotically asynapsis-induced aneuploidy in autopolyploid Arabidopsis thaliana. J Plant Res 123:87–95

19. Ferdous M, Higgins JD, Osman K, et al (2012) Inter-homolog crossing-over and synapsis in Arabidopsis meiosis are dependent on the chromosome axis protein AtASY3. PLoS Genet 8:e1002507

20. Pochon G, Henry IM, Yang C, et al (2022) The Arabidopsis Hop1 homolog ASY1 mediates cross-over assurance and interference. bioRxiv 2022.03.17.484635

21. Kloosterman WP, Guryev V, van Roosmalen M, et al (2011) Chromothripsis as a mechanism driving complex de novo structural rearrangements in the germline. Hum Mol Genet 20:1916–1924

22. Zhang C-Z, Spektor A, Cornils H, Francis JM, Jackson EK, Liu S, Meyerson M, Pellman D (2015) Chromothripsis from DNA damage in micronuclei. Nature 522:179–184

23. Ly P, Brunner SF, Shoshani O, et al (2019) Chromosome segregation errors generate a diverse spectrum of simple and complex genomic rearrangements. Nat Genet 51:705–715

24. Kneissig M, Keuper K, de Pagter MS, et al (2019) Micronuclei-based model system reveals functional consequences of chromothripsis in human cells. Elife. https://doi.org/10.7554/eLife.50292

25. Umbreit NT, Zhang CZ, Lynch LD, et al (2020) Mechanisms generating cancer genome complexity from a single cell division error. Science. https://doi.org/10.1126/science.aba0712

26. Papathanasiou S, Markoulaki S, Blaine LJ, Leibowitz ML, Zhang C-Z, Jaenisch R, Pellman D (2021) Whole chromosome loss and genomic instability in mouse embryos after CRISPR-Cas9 genome editing. Nat Commun 12:5855

27. Chiang C, Jacobsen JC, Ernst C, et al (2012) Complex reorganization and predominant non-homologous repair following chromosomal breakage in karyotypically balanced germline rearrangements and transgenic integration. Nat Genet 44:390–397

28. Pellestor F, Gatinois V, Puechberty J, Geneviève D, Lefort G (2014) Chromothripsis: potential origin in gametogenesis and preimplantation cell divisions. A review. Fertil Steril 102:1785–1796

29. Cuacos M, Lambing C, Pachon-Penalba M, Osman K, Armstrong SJ, Henderson IR, Sanchez-Moran E, Franklin FCH, Heckmann S (2021) Meiotic chromosome axis remodelling is critical for meiotic recombination in Brassica rapa. J Exp Bot 72:3012–3027

30. Yuan W, Li X, Chang Y, Wen R, Chen G, Zhang Q, Wu C (2009) Mutation of the rice gene PAIR3 results in lack of bivalent formation in meiosis. Plant J 59:303–315

31. Golubovskaya I, Avalkina NA, Sheridan WF (1992) Effects of several meiotic mutations on female meiosis in maize. Dev Genet 13:411–424

32. Cao L, Wang S, Zhao L, Qin Y, Wang H, Cheng Y (2021) The Inactivation of Arabidopsis UBC22 Results in Abnormal Chromosome Segregation in Female Meiosis, but Not in Male Meiosis. Plants. https://doi.org/10.3390/plants10112418

33. Henry IM, Zinkgraf MS, Groover AT, Comaia L (2015) A system for dosage-based functional genomics in poplar. Plant Cell 27:2370–2383

34. Li H, Durbin R (2009) Fast and accurate short read alignment with Burrows–Wheeler transform. Bioinformatics 25:1754–1760

